# BioViz *Connect*: Web application linking CyVerse cloud resources to genomic visualization in the Integrated Genome Browser

**DOI:** 10.1101/2020.05.15.098533

**Authors:** Karthik Raveendran, Nowlan H Freese, Chaitanya Kintali, Srishti Tiwari, Pawan Bole, Chester Dias, Ann E Loraine

## Abstract

Genomics researchers do better work when they can interactively explore and visualize data. However, due to the vast size of experimental datasets, researchers are increasingly using powerful, cloud-based systems to process and analyze data. These remote systems, called science gateways, offer user-friendly, Web-based access to high performance computing and storage resources, but typically lack interactive visualization capability. In this paper, we present BioViz *Connect*, a middleware Web application that links the CyVerse science gateway to the Integrated Genome Browser (IGB), a highly interactive native application implemented in Java that runs on the user’s personal computer. Using BioViz *Connect*, users can (i) stream data from the CyVerse data store into IGB for visualization, (ii) improve the IGB user experience for themselves and others by adding IGB specific metadata to CyVerse data files, including genome version and track appearance, and (iii) run compute-intensive visual analytics functions on CyVerse infrastructure to create new datasets for visualization in IGB or other applications. To demonstrate how BioViz *Connect* facilitates interactive data visualization, we describe an example RNA-Seq data analysis investigating how heat and desiccation stresses affect gene expression in the model plant *Arabidopsis thaliana*. Lastly, we discuss limitations of the technologies used and suggest opportunities for future work. BioViz *Connect* is available from https://bioviz.org.

## Introduction

Science gateways are Web sites that implement user-friendly interfaces to high performance computing and storage systems (Wilkins-Diehr et al., 2008). Science gateways typically assemble and curate discipline-specific, command-line, Unix-based tools within a single, easy-to-use interface, enabling users to run compute-intensive processing on datasets too large for a personal computer (Giardine et al., 2005; Goff et al., 2011; Merchant et al., 2016). In a typical use case, domain researchers upload their “raw” (unprocessed) data to the gateway site and then operate the gateway’s Web-based interface to create custom processing and analysis pipelines, where a pipeline is defined as tasks performed in sequence by non-interactive tools which emit and consume well-understood file types and formats. Common pipeline tasks in genomics include aligning RNA-Seq sequences onto a reference genome to produce BAM (binary alignment) format files (Li et al., 2009), generating scaled RNA-Seq coverage graphs from the “BAM” files using tools such as genomeCov (Ramirez et al., 2016), or searching promoter regions for sequence motifs common to sets of similarly regulated genes using tools such as DREME (Bailey, 2011).

A science gateway aims to provide a single point of access for tools needed to process and analyze data from a research project. However, native visualization tools with their own graphical user interfaces separate from a Web browser are difficult to use with Web-based science gateway systems. The Integrated Genome Browser from BioViz.org (Nicol et al., 2009; Freese et al., 2016) and the Broad Institute’s Integrative Genomics Viewer (Robinson et al., 2011) exemplify this problem. Both tools require that data files reside on the user’s local file system or that they be accessible via HTTP (hypertext transfer protocol) and addressable via a file-specific URL (Uniform Resource Locator). If the gateway system does not allow URL-based access to data, then users must download the data files onto their local computer file system, which may not be practical or allowed.

Related problems confront visualization systems implemented as Web applications, deployed on Web hosts and not the user’s local computer. Using Web applications to visualize data can be even more challenging for users, because these applications often require hard-to-set-up data storage and delivery mechanisms specialized to the application. To view one’s data using the Web-based UCSC Genome Browser software, users can either deploy their own copy of the software, which is difficult, or they can instead set up a UCSC Track Hub server, which is less technically challenging but nonetheless requires a complex set of files to be created and configured (Raney et al., 2014). Similarly, using the JBrowse Web-based genome browser requires deploying data in JBrowse-compatible formats (Buels et al., 2016).

Another typical requirement for science gateways is extensibility, meaning they require a way for gateway developers or users to add new tools to the system to accommodate or even potentiate new directions for research. The CyVerse science gateway, the focus of this article, supports extensibility by allowing developers to create and deploy CyVerse Apps, which are user-contributed container images that run within a CyVerse-provided container environment (Devisetty et al., 2016). Users create containers using Docker and then contribute their container image along with metadata specifying input parameters and accepted data types to CyVerse. Once accepted and deployed, the container is configured to run as an asynchronous “job” within the CyVerse infrastructure via a queuing system. Thus, Apps run non-interactively and therefore are not well-suited to providing interactive, exploratory visualization. However, these Apps do provide a means to create new input data for visualization, as we explore here.

In this paper, we introduce BioViz *Connect*, a Web application that overcomes limitations described above to add genome visualization capability to the CyVerse Discovery Environment, part of the CyVerse science gateway system. Previously called iPlant, the CyVerse science gateway is a United States National Science Foundation funded cyberinfrastructure project with the aim of providing computational resources for life sciences researchers (Goff et al., 2011; Merchant et al., 2016). We chose to work with the CyVerse Discovery Environment in this study because it features a rich Application Programming Interface (API), the Terrain REST API, that supports secure computational access to CyVerse data storage and analysis resources. Using this API, we implemented a new visualization-focused interface to these resources, called BioViz *Connect*, using the Integrated Genome Browser (IGB) as the demonstration application. We selected IGB because it offers one of the richest feature sets for visual analysis in genomics (for descriptions of IGB functionality, see (Nicol et al., 2009; Gulledge et al., 2014; Loraine et al., 2015; Freese et al., 2016; Mall et al., 2016)) and because we are members of the core IGB development team. Therefore, we possessed insider’s knowledge of the featured visualization application that allowed us to modify IGB as needed for the project.

BioViz *Connect* enables users of Integrated Genome Browser to visually analyze their CyVerse data without having to download entire files to their local computer or migrate their data into application specific data stores. In the following sections, we describe how BioViz *Connect* is implemented, explaining the technology stack used and how BioViz *Connect* interacts with the CyVerse science gateway resources via its Terrain API. Next, we describe how BioViz *Connect* enables flow of data into the IGB desktop software by activating a REST API endpoint residing in IGB itself. To illustrate the functionality, we describe an example use case scenario for BioViz *Connect* in which a hypothetical analyst uses visualization and visual analytics tools within IGB in conjunction with their CyVerse account to quality-check and analyze an RNA-Seq data set from *Arabidopsis thaliana* plants undergoing desiccation and heat stresses. Lastly, we discuss insights gained from implementing BioViz *Connect*, describe limitations of the technology used, and propose how these limitations might be overcome. BioViz Connect represents a next step toward building integrated, user-friendly computational environments that blend powerful local tools like IGB with even more powerful remote infrastructures like CyVerse, creating new possibilities for users to discover biologically meaningful features in data while avoiding artifacts.

## Materials and Methods

### BioViz *Connect* client and server-side design

BioViz *Connect* consists of two parts: a JavaScript-based user interface that runs in a Web browser and a server-side application that manages authentication and communication with Terrain API endpoints. The user interface code on the client-side is implemented using HTML5, CSS, Bootstrap 4.3.1, JavaScript, and jQuery 1.10.2. The server-side code is implemented in python3 using the Django web application framework (Fig. 1). The currently available production instance of BioViz *Connect* is deployed on an Ubuntu 18.04 system and hosted using the apache2 Web server software as a reverse proxy. BioViz *Connect* code is open source and available from https://bitbucket.org/lorainelab/bioviz-connect.

**Figure 1.**
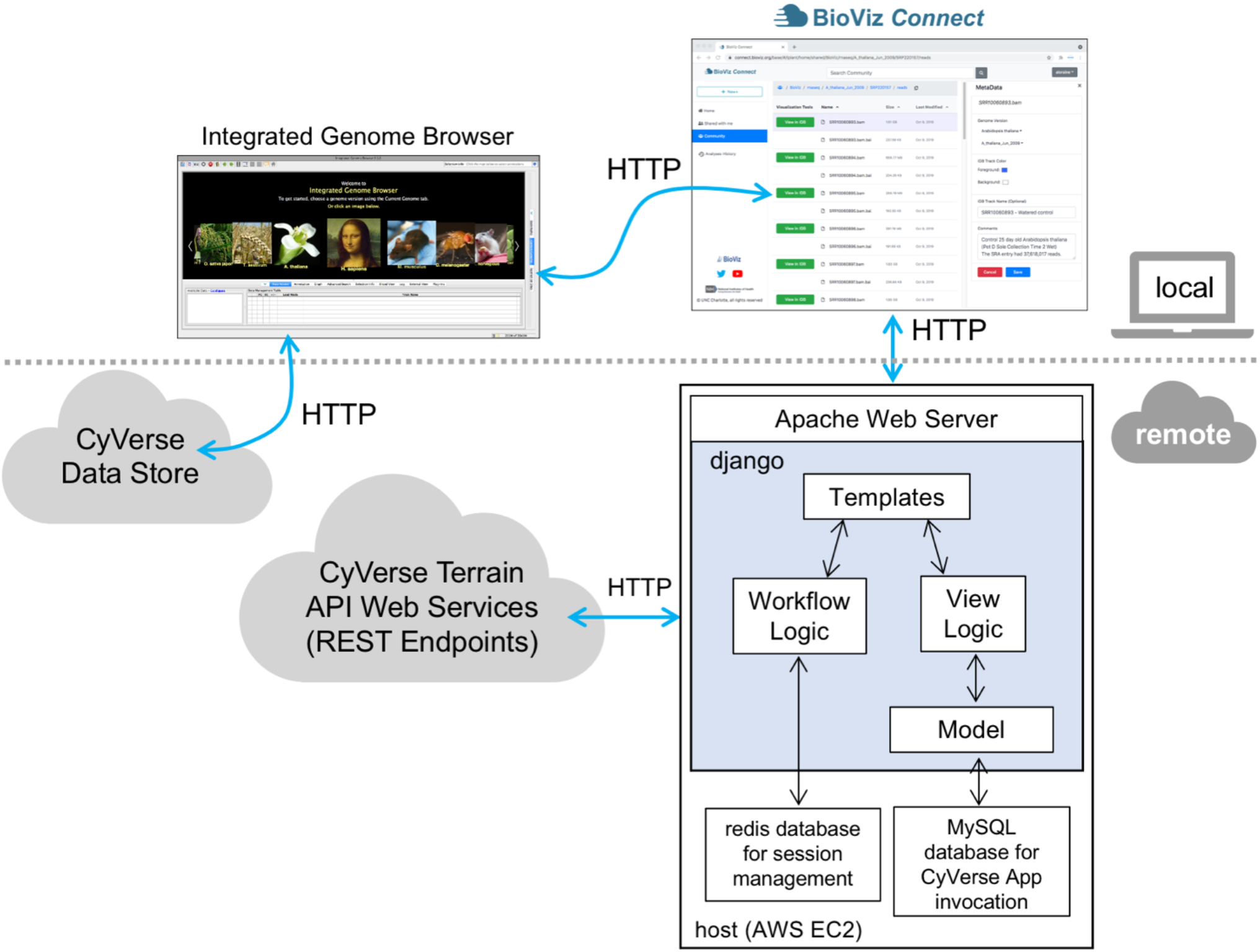
Diagram illustrating local client and remote server-side design of BioViz *Connect*.

### User and password management

To use BioViz *Connect*, users must first obtain a CyVerse Discovery Environment account by registering at https://user.cyverse.org/register (CyVerse, RRID:SCR_014531). At the time of this writing, there is no charge for this account. BioViz *Connect* delegates user management, including logging in and password management, to the Central Authentication Service (CAS) OAuth service hosted and maintained by CyVerse (Fig. 1). Thus, CyVerse infrastructure manages user accounts and information; no BioViz-specific accounts or passwords are required. After a user has logged in to BioViz Connect using their CyVerse credentials, BioViz *Connect* uses a server-side Redis database to store a user-specific access token for the duration of the session, allowing it to access and modify user data stored in CyVerse via the Terrain API on the user’s behalf.

### How BioViz *Connect* communicates with Integrated Genome Browser

Integrated Genome Browser is a free, open-source desktop software program written in Java which users download and install on their local computer systems (IGB, RRID:SCR_011792) (Nicol et al., 2009; Freese et al., 2016). Installers for Linux, MacOS, and Windows platforms are available at https://bioviz.org. The IGB source code resides in a git repository hosted on Atlassian’s bitbucket.org site (https://bitbucket.org/lorainelab/integrated-genome-browser). IGB version 9.1.4 or greater is required for IGB to connect to BioViz *Connect*.

IGB contains a simple Web server configured to respond to REST-style queries on an IGB-specific port on the user’s local computer. Javascript code downloaded into the Web browser when users visit BioViz *Connect* pages enables requesting URLs addressed to “localhost”, the user’s computer, using the IGB-specific port. IGB intercepts these requests and performs actions based on URL parameters.

### BioViz *Connect* metadata

BioViz *Connect* uses the Terrain Metadata API to manage and obtain IGB-specific metadata for files and folders. The Terrain API represents metadata items as triplets containing Attribute, Value, and Unit. A metadata item’s Attribute attaches meaning to what the metadata contains, and application developers can create their own custom Attributes to support diverse purposes. For example, since BioViz *Connect* is concerned with genomic data visualization, we created custom Attributes signaling genome assembly version, visual style information such as foreground color and background color, and free text comments on the data provided by the user, which are displayed in BioViz *Connect*’s Web interface. A metadata item’s Value is specific to the file or folder being tagged. BioViz *Connect* uses the Unit value to indicate that the metadata element concerns IGB and the BioViz Connect application.

The genome identifier attribute requires further explanation, as matching genome version names across systems has been a major source of problems for users. Integrated Genome Browser, like many other systems, uses an application-specific scheme for naming genome versions, and contains a listing of synonyms matching these IGB-specific names onto genome version names from other systems. For example, the IGB genome version named H_sapiens_Feb_2009 is the same as UCSC genome version name hg17, which is the same as NCBI version 35. The BioViz *Connect* user interface includes components for users to view, designate, or change the genome version metadata associated with individual files. To ensure compatibility with IGB, BioViz *Connect* uses a list of IGB-formatted genome identifiers hosted on the IGB Quickload site (http://igbquickload.org/quickload/) to configure the genome version selection components, implemented as menus. When users operate the interface to view data within IGB, the genome version metadata, along with style metadata, are passed to IGB via its localhost REST endpoint. This ensures that the data appear in the context of the correct genome assembly, alongside other data already loaded from BioViz *Connect* or other sources, while also enabling the user to specify in advance how the data will look once it appears in IGB. In addition, if other users load the same files, the data will look the same.

### Enabling access to data via public URLs

The flow of data from CyVerse into IGB depends on two key technical features of the CyVerse data storage and hosting system. The CyVerse API enables users to create publicly accessible URLs for data files in their accounts, and these URLs can be created or destroyed at will. In the current implementation, URLs created in this way are accessible to any internet user. Second, the CyVerse infrastructure supports HTTP range requests for these URLs, enabling client such as IGB to request subsets of data, thus avoiding having to download an entire data file.

The BioViz *Connect* interface is designed to make the process of managing these URLs as easy as possible, similar to commercial cloud storage systems such as Dropbox and Google Drive that let users create, destroy, and manage public links to individual files and folders. Within the BioViz *Connect* interface, users create URLs for individual files by right clicking the file and selecting the “Manage Link” option (Fig. 2).

**Figure 2.**
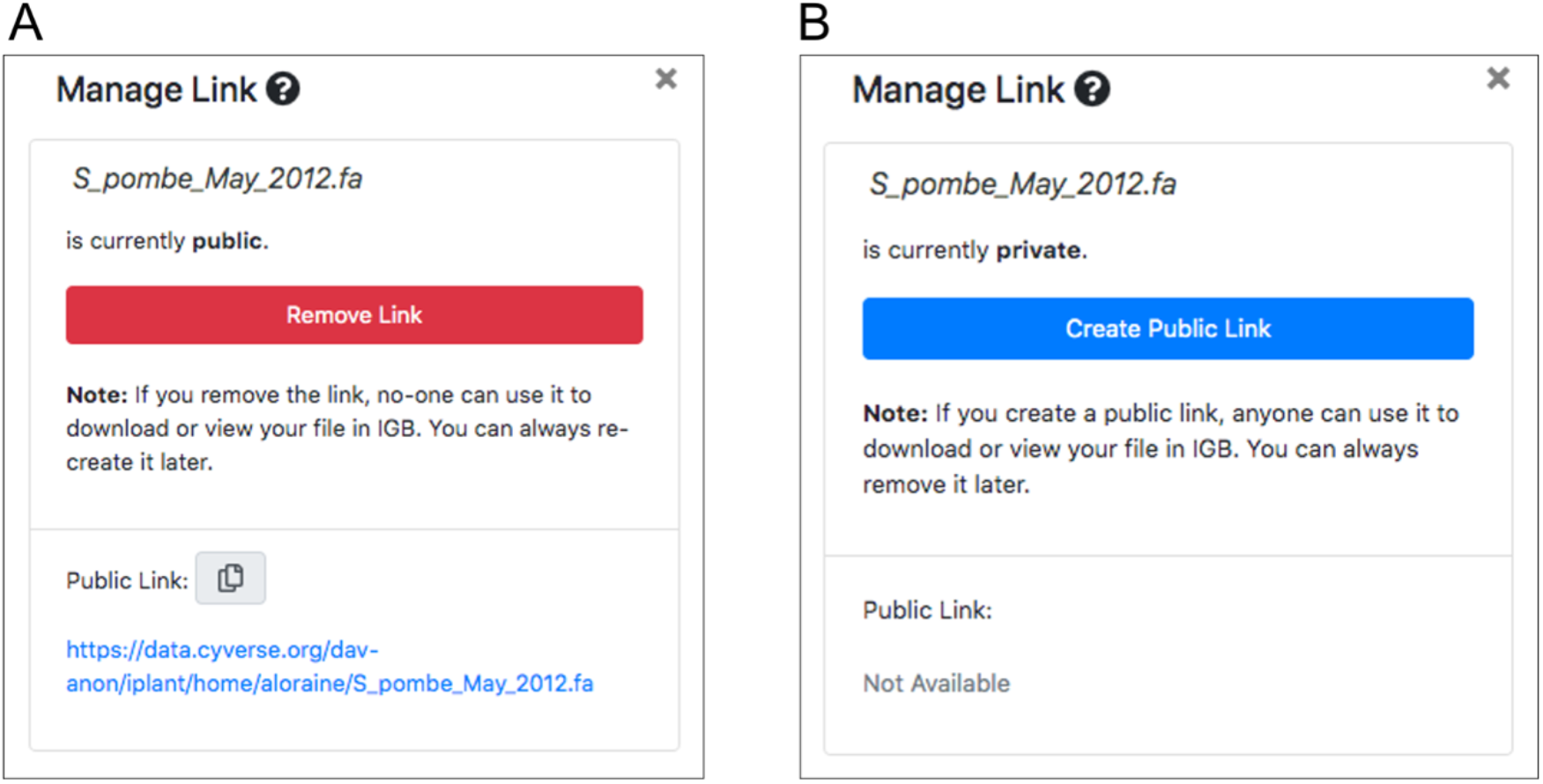
BioViz *Connect* “Manage Link” interface, from the right panel display. (A) Interface when a file has a publicly accessible link. (B) Interface when the file has no public link.

### BioViz *Connect* deployment

BioViz *Connect* is managed using ansible roles and playbooks publicly available in a git repository from https://bitbucket.org/lorainelab/bioviz-connect-playbooks. The playbooks contain two sets of tasks. One set of tasks creates a virtual machine using the Amazon EC2 Web service. Once the host is created and running, a second set of ansible tasks installs and configures software on the host, including an Apache2 Web server, a MySQL database, and the BioViz *Connect* code base. Playbook users can specify the BioViz *Connect* repository and branch they wish to deploy, which facilitates rapid testing of proposed new code. During the provisioning process, a call is made to a Terrain endpoint that provides a list of all CyVerse asynchronous analysis apps that can produce output visible to IGB. These data are then used to construct the “analysis” sections of the user interface.

### BioViz *Connect* interface for running visual analysis Apps

When users right-click a file name in BioViz *Connect*, a context menu appears with an option labeled “Analyse.” Information about IGB-compatible Apps, the file types they can accept, and App parameters are stored in a relational database. When a user selects this option, BioViz *Connect* queries the database to identify IGB Community Apps that accept the file as input, and these are then displayed to the user. Once the user has selected an App, another query retrieves additional information about it, such as user-friendly description of what the App does, which is then displayed to the user.

The CyVerse ecosystem contains many hundreds of Apps, many of which are redundant or obsolete, and so the BioViz Team controls which ones are shown to users by adding them to the IGB Community, a CyVerse organizing concept that groups resources (such as Apps) according to which users can use or modify them. BioViz *Connect* only shows Apps that have been added to the IGB Community.

### RNA-Seq data

RNA-Seq data presented in the use case scenario are from Sequence Read Archive Bioproject PRJNA509437 (Leinonen et al., 2011), an experiment in which Arabidopsis plants underwent either a 3-hour, non-lethal heat stress or a multi-day desiccation stress. Two post-treatment sample time points were collected for treated plants and their untreated control counterparts, with two to four replicates per sample type and 23 samples in total. Sample libraries were sequenced in single-end runs of the Illumina platform and are identified by their run identifiers. BAM files were generated by aligning sequence reads to the Arabidopsis June 2009 reference genome assembly using TopHat2 (TopHat, RRID:SCR_013035) (Kim et al., 2013). The data are available in the Community folder of publicly accessible datasets, represented as a folder in the left-side panel of the BioViz *Connect* display.

## Results

### Understanding and navigating the BioViz *Connect* interface

To get started using BioViz Connect, the user opens the BioViz.org website in a Web browser, selects the link labeled BioViz *Connect*, and then clicks the link labeled “Sign in with your CyVerse ID”. This action opens a Central Authentication Service (CAS) page, hosted by CyVerse, where users enter their CyVerse username and password, or sign up for a new account if they do not already have one. Once the user has logged in, the Web browser then returns to a “call-back” URL on the BioViz.org site, which displays the BioViz *Connect* user interface, a browsable, sortable, paginated view of the user’s CyVerse home directory and its contents (Fig. 3A).

**Figure 3.**
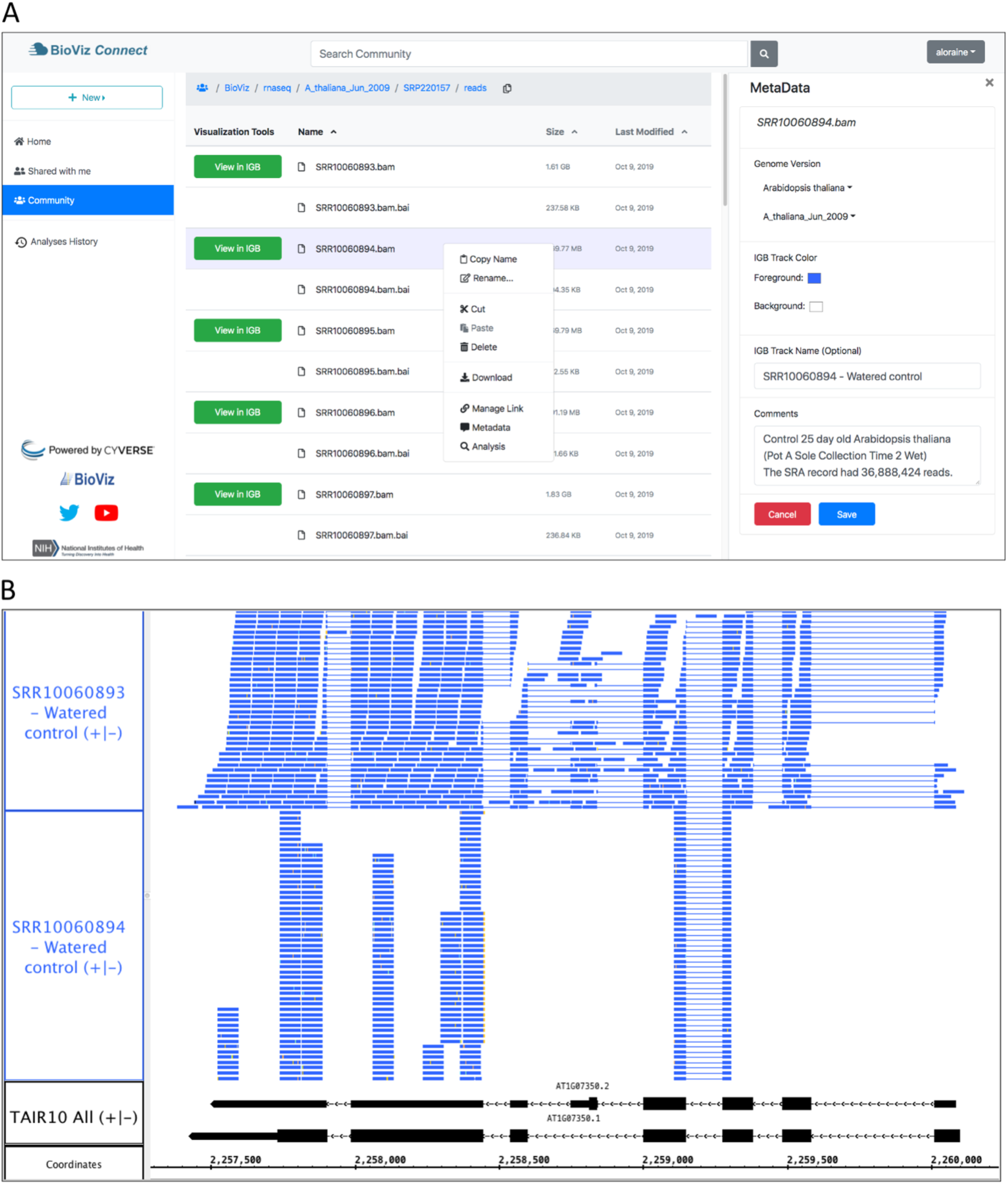
Example BioViz *Connect* main page and data visualization. (A) BioViz *Connect* main page. The left panel shows shortcuts to home, shared, community folders. The middle panel lists files and folders. The right panel shows the selected file’s metadata. (B) SRR10060893.bam and SRR10060894.bam files viewed in IGB overlapping the SR45a gene of *Arabidopsis thaliana*. The track labeled TAIR10 mRNA shows SR45a gene models AT1G07350.1 and AT1G07350.2.

This view of files and data resembles the interface for commercial, consumer-focused cloud storage systems, a deliberate design choice aimed at building on many users’ familiarity with Google Drive, the Dropbox Web interface, and others. This interface displays a sortable, table-based view of the user’s home directory within the CyVerse file storage system, displaying a listing of files and folders the user has uploaded to their account or created using CyVerse Apps, including BioViz *Connect* Apps described in later sections. Single-clicking a file or folder selects it, double-clicking a folder opens it and displays the contents, and double-clicking a file opens a metadata display showing information about the file. A bread crumb display at the top of the page shows the path from the root folder to the currently opened folder, and a copy icon next to the breadcrumb allows the user to copy the folder name and path. The browser forward and back buttons work as expected, and users can bookmark individual screens for faster navigation. The URLs displayed in the browser’s URL bar match the currently opened folder’s location, making the interface feel more polished and user-friendly by ensuring that every user-facing detail, including the URL, mimic and reinforce how the user has organized their data within the CyVerse virtual file system.

The top part of every BioViz *Connect* page also features a search bar that can be used to find files and folders with names matching a user-entered query string. Matches are returned in a list view similar to the original table view, and users can sort the results list by name, size, or date modified. Only files for which the user has read access and that reside in the currently visible section (Home, Community, or Shared with me) are returned. On the left side of every page, BioViz *Connect* displays icons representing shortcut links to the user’s home directory, a publicly available community data folder, and other destinations. The “Community” folder contains data published for all CyVerse users, including the example RNA-Seq data set for the use case scenario described in the next section.

### Using BioViz *Connect* to view data in Integrated Genome Browser

To demonstrate BioViz *Connect* functionality, we next describe an example use case scenario in which a hypothetical researcher visually analyzes data from a typical RNA-Seq experiment. The use case focuses on two main tasks: visually checking data quality and then confirming differential expression of a control gene known to be regulated by the treatment. Data files discussed below are available in the Community folder in file path “*BioViz / rnaseq / A_thaliana_Jun_2009 / SRP220157 / reads”*.

The experimental design included two treatments, heat and desiccation stress, their controls, and two time points, totaling six sample types, each with two to four replicates. The RNA-Seq sequences are available in the Sequence Read Archive, and the researcher has obtained the data, aligned it to the reference genome, and then contributed the files to the Community folder. Alignment files are stored in the file path *“BioViz / rnaseq / A_thaliana_Jun_2009 / SRP220157 / reads”*. The user has also annotated each file using the BioViz Connect interface, adding the genome version, visual style information, and notes describing each sample.

Now that the data are organized and annotated, the researcher uses the BioViz *Connect* interface to import the data into Integrated Genome Browser for visualization and proceeds to look at each file, one by one, to check the quality of the alignments and confirm file identity. BioViz *Connect* makes this task easy to perform. To illustrate, we discuss RNA-Seq alignment files SRR10060893.bam and SRR10060894.bam, replicate control samples from time point one of the heat stress treatment. A quick scan of files listed in the BioViz *Connect* table view shows that SRR10060893.bam has size 1.61 GB, about twice the size of SRR10060894.bam, which is 0.669 GB. The user has annotated the files with the number of sequence reads obtained per sample, around 37 million for each. Because the samples were sequenced to about the same depth, their resulting alignment files ought to have similar sizes. Visualizing the sequence read alignments will help explain the discrepancy.

To visualize the alignments, the user launches Integrated Genome Browser, which is already installed on the local computer, downloaded from the BioViz.org Web site. Once IGB is running, the user clicks the “View in IGB” button available in the “Visualization Tools” column in the BioViz *Connect* table view, repeating this action for each file. This action causes Javascript code running within the Web browser to request data from a local URL (domain “localhost”) corresponding to a REST endpoint implemented within IGB. The URL includes parameters such as the publicly accessible URL for the data file, the IGB name of its reference genome, and visual style information indicating how the file should look once loaded into IGB. In response, IGB opens the requested genome version associated with the file and adds the file as new tracks to the display.

To check assumptions about a new data set, it is useful to visualize a gene of known behavior, such as a gene already known to be regulated by the experimental treatment. Prior work from our lab and others have shown that SR45a, encoding an RNA-binding protein, is up-regulated by heat and desiccation stresses, making it a good choice for this purpose (Yoshimura et al., 2011; Gulledge et al., 2012). To find the gene, the analyst enters SR45a into IGB’s search interface at the top left of the IGB window, which zooms and pans the display to the gene’s position in the genome. Next, the user loads the alignments into the display by clicking the “Load Data” button at the top right of the IGB window. Once the data load, the user customizes track appearance by modifying vertical zoom setting and changing the number of sequences that can be shown individually in a track (stack height), creating the view shown in Figure 3B.

This customized view makes problems with SRR10060894 obvious at a glance. The alignments for this sample appear to stack on top of each other in orderly, uniform towers covering only 30% of the gene’s exonic sequence. By contrast, the alignments for sample SRR10060893 cover most of the exonic sequence and also include many spliced reads split across introns. The sparser pattern observed in SRR10060894 typically arises when the library synthesis process included too many polymerase chain reaction amplification cycles, reducing the diversity of resulting sequence data. This pattern indicates that the user should exclude SRR10060894 from further analysis, but the other file appears to be fine.

### Comparing sequencing depth and complexity using IGB visual analytics

Repeating the preceding process with other samples in the dataset, the user identifies another problematic pair of files. The files are replicates, but like the previous example, the files sizes differ. The alignments file SRR10060911.bam is 1.83 Gb, but its replicate SRR10060912.bam is only 0.454 Gb. Opening and viewing the alignment files in IGB, the user confirms that one file appears to contain more data than the other (Fig. 4A). To quantify this observation, the user takes advantage of a simple, interactive visual analytics feature within IGB: selection-based counting. As with PowerPoint and many other graphical applications, IGB users can click-drag the mouse over graphical elements to select a group of items and then single-click while pressing SHIFT or CTRL-SHIFT keys to add or remove items from selection group. IGB reports the number of currently selected items in the Selection Info box at the top right of the IGB window. Using this feature, the researcher finds that sample SRR10060912 contains 1,925 alignments covering SR45a, and sample SRR10060911 has 10,867 alignments, nearly five times as many.

**Figure 4.**
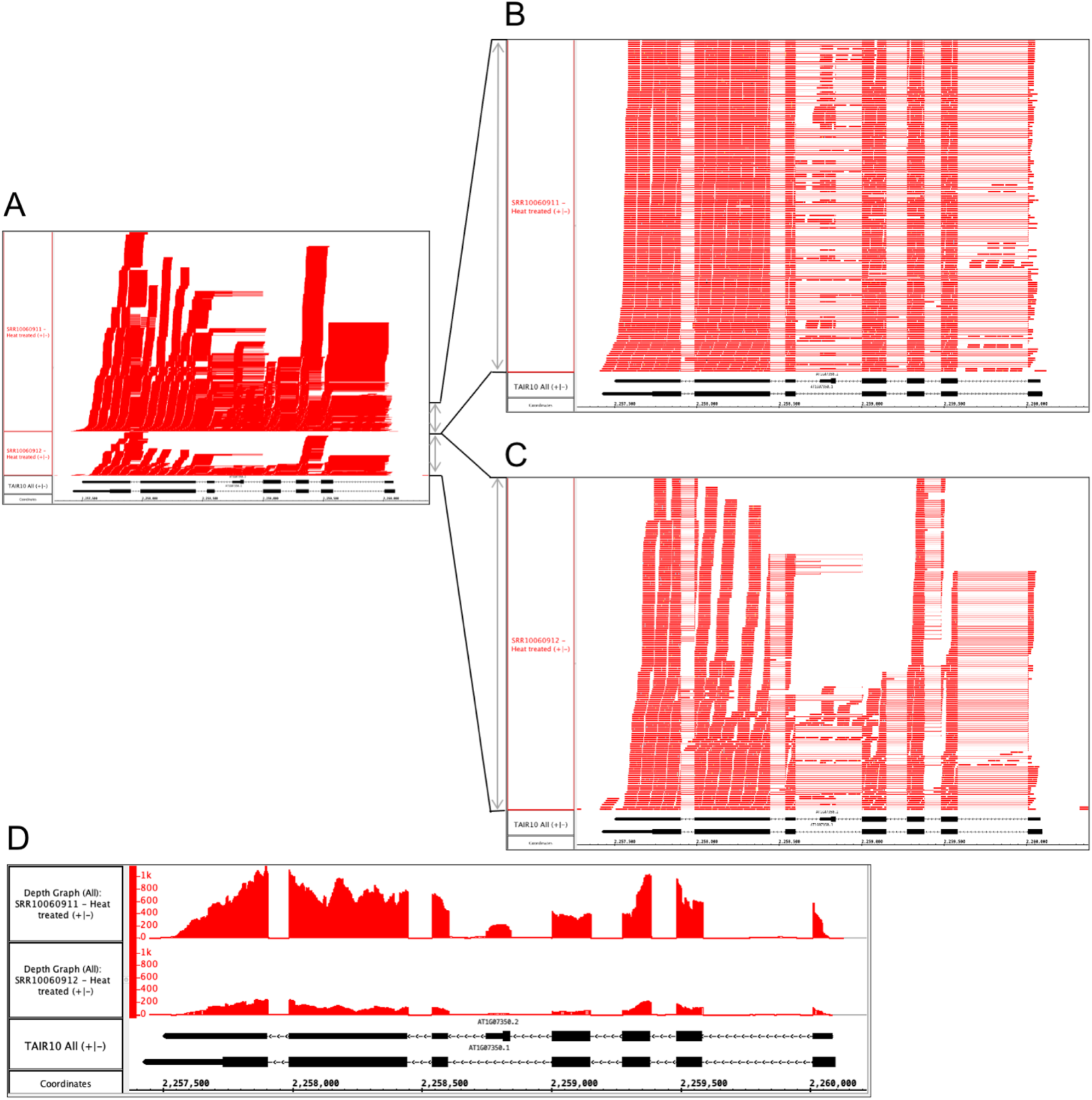
Heat treated samples viewed in IGB. (A) Vertical dimension is compressed to show all alignments. (B) SR10060912 and (C) SR10060911 tracks stretched vertically to reveal alignment patterns in more detail. (D) Alignment coverage graphs calculated within IGB using alignments from A. The y-axis values represent the number of aligned sequences per base pair position indicated on the coordinates track. The track labeled TAIR10 mRNA shows SR45a gene models AT1G07350.1 and AT1G07350.2.

By further configuring track height and appearance settings, and operating IGB’s dynamic vertical and horizontal zoom controls, the user can stretch the display in each dimension independently to reveal more detail about the alignments (Fig. 4B and 4C). From this new view of the data, the user can tentatively conclude that alignment pattern diversity is similar in each sample, but the depth of sequencing was greater in SRR1006911. To confirm the finding, the user then applies a visual analytics function (called a “Track Operation” within IGB) that creates coverage graphs, also called depth graphs, using data from the read alignment tracks (Fig. 4D). To make a coverage graph, the user right-clicks a track label for a read alignment track and chooses option “Track Operations > Depth Graph (All)”. This generates a new track showing a graph in which the y-axis indicates the number of sequences aligned per x-axis position, corresponding to base pair positions. After modifying the y-axis lower and upper boundary values (using controls in IGB’s Graph tab), the user again can observe that the pattern of alignments is similar between the two samples, but the overall level of sequencing was different. Thus, the file size difference most likely is due to a difference in sequencing depth rather than a problem with the library synthesis, as was the case in the previous example.

### Normalizing coverage graphs to compare gene expression visually

Coverage graphs set to the same scale allow comparing gene expression across sample types, but only if the libraries were sequenced to approximately the same depth. If not, then coverage graphs need to be normalized before comparing them. Scaling coverage graphs within IGB is impractical, however, as it would require downloading, reading, and processing the entire bam-format alignments file. A better approach is to off-load computationally intensive visual analytics tasks to CyVerse cloud computing resources. To demonstrate the value of this strategy, we deployed the deepTools genomeCoverage command line tool from the deepTools suite (Deeptools, RRID:SCR_016366) as a new IGB-friendly CyVerse App (Ramirez et al., 2016).

To create a scaled coverage graph, the user returns to BioViz *Connect*, right-clicks a bam format file, and chooses “Analyse.” This opens the Analysis right-panel display, which lists all IGB-compatible CyVerse Apps that can accept the selected file type as input (Fig. 5A). Selecting “Make scaled coverage graph” opens a form with options for creating the graph using the genomeCoverage algorithm (Fig. 5B). The interface includes a place for the user to enter names for the analysis and for the output file that will be produced. The user then clicks “Run Analysis” button, which calls upon the CyVerse analysis API to run the App with specified parameters using CyVerse computing resources. The request to run the App and the work it performs are called “jobs,” and jobs are carried out asynchronously, running and completing only when resources they require become available, as with other systems set up for high-performance computing. Users can check job status by using the Analyses History in the BioViz Connect interface (Fig. 5C), where Analyses are listed as Queued (waiting to run), Running, Failed, or Completed. Independent of BioViz *Connect*, the CyVerse infrastructure sends an email to users when jobs finish. When a job finishes, any files or folders it creates appear in the analyses folder in the user’s home directory, or in the same location as the input files, if those are stored in a location where the user has permission to modify or add to the folder. To quickly navigate to results, users can click the analysis name in the Analyses History, opening the folder where the output data files are stored.

**Figure 5.**
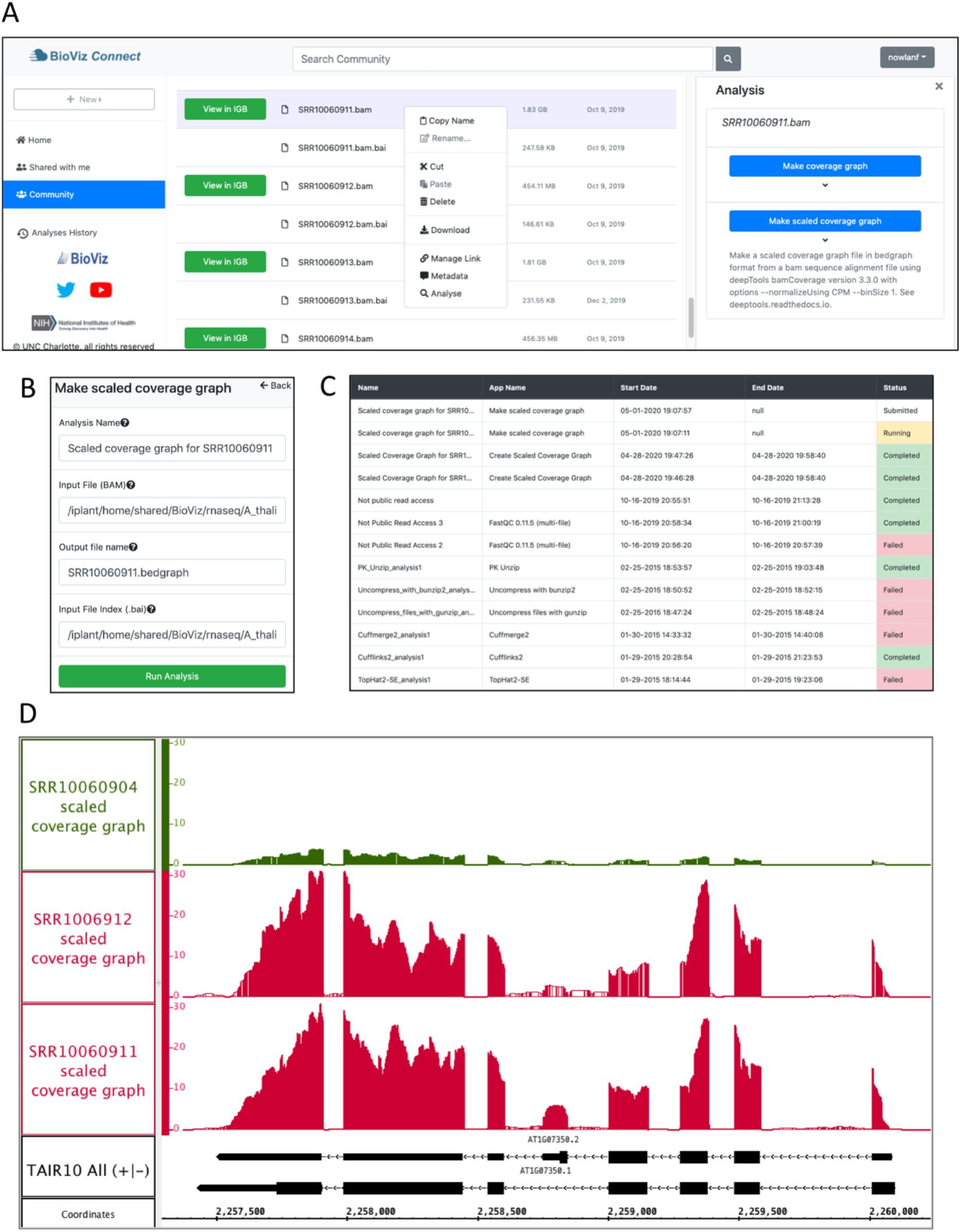
Example analysis in BioViz *Connect* with output visualized in IGB. (A) BioViz *Connect* main page with analysis right panel open. (B) Scaled coverage graph analysis options for naming the analysis, selecting input file, output file name, and index file selection. (C) Analyses History showing the status of current and previous jobs. (D) SRR10060902 (control), SRR10060911 (heat treated), and SRR10060912 (heat treated) scaled coverage graphs viewed in IGB overlapping the SR45a gene of *Arabidopsis thaliana*.

Figure 5D shows sample App output, a visualization of the SR45A region with three scaled coverage graphs loaded from bigwig data files, a compact binary format for representing numeric values associated with base pairs in a genome map. Two heat-treated and one control sample are shown. The tree coverage graphs have been configured to use the same y-axis scale, making it obvious that the heat treatment elevated SR45A gene expression, consistent with previously published reports. The image presents a clear visual argument in favor of this conclusion, and it also shows the user how much the expression level measurement varies across the gene body, something a single summary statistic cannot provide.

## Discussion

BioViz *Connect* introduces and demonstrates innovations in the field of science gateway development and research, while providing useful functionality for researchers seeking to understand and visualize genomic data. BioViz *Connect* enables users of the CyVerse science gateway to visualize genomic data files from their CyVerse accounts using Integrated Genome Browser, a desktop application. To our knowledge, BioViz *Connect* is the first and only resource that integrates remote CyVerse file storage and computational resources with a genome browser native to the local computer, achieving this cross-application communication via localhost REST endpoints.

We implemented BioViz *Connect* using the CyVerse Terrain API, a collection of remote REST endpoints that form a comprehensive computational interface to CyVerse resources. BioViz *Connect* is the first application developed using the Terrain API by a group outside the CyVerse development team. Because our work is open source, developed entirely in public, other groups can use our implementation as a guide or inspiration for their own work. BioViz *Connect* further demonstrates to the larger community of biologists, developers, and funders that modern, feature-rich REST interfaces to powerful computational resources stimulate and enable innovation and progress.

The scaled coverage graphs described in the use case scenario offer a useful, practical example of how remote resources can power interactive visual analytics on the desktop, an idea that has been explored in diverse fields and settings, but not often applied to genome visualization as was done here. The example we presented used a pre-existing algorithm, developed by others, but it shows how developers can harness a more powerful gateway system to develop and deploy all-new interactive genome data visualizations. Offloading compute-intensive visual analytics functions to science gateway systems like CyVerse will likely become more appealing and important as the size and complexity of genomic data continue to increase.

However, at least two important technical limitations remain, providing opportunities for future work. The first technical limitation has to do with how data flow from the CyVerse back end data store and into the desktop genome browser application. Integrated Genome Browser as currently implemented can display CyVerse data because the CyVerse API can assign publicly accessible URLs to individual data files, which makes them available for visualization but exposes them to everyone on the internet. This problem of public accessibility could perhaps be addressed by adding password protection to these URLs, using Basic Authentication headers defined by the HTTP protocol. IGB already supports logging into password-protected Web servers, and so this solution would require little or no changes on the client side.

A more subtle problem has to do with the data file formats themselves. IGB, along with every other genome visualization system we are aware of, uses random access, indexed file formats to retrieve subsets of data corresponding to genomic regions. For example, BAM (binary alignment) files are typically large, impractical to download in their entirety. The data stored in these files are sorted by genomic location and therefore can be indexed by genomic location. When retrieving data for a desired genomic region, IGB and other programs use the index to look up the range of bytes where those data reside in the target file, and then read and process only the data for that region, ignoring the rest. This idea of mapping genomic coordinates to physical file coordinates has been in heavy use for decades, for as long as IGB has existed. Indeed, the original IGB development team at Affymetrix implemented one of the first indexed file formats, called “bar” for “binary array format”, used for storing and accessing data from Affymetrix genome tiling arrays, one of the first technologies invented to survey transcription across an entire genome in an unbiased way. However, this method of data access tends to expose information to the client visualization software in ways that might not be desirable for some use cases. For example, a physician may want to view the results of sequencing a particular gene for a patient, and to do so, the visualization application needs access to the index in order to request the required subset of data. As we and others have explored (Pedersen et al., 2017; Loraine et al., 2021), the index itself contains a visualizable map of an individual’s genome.

The second technical limitation concerns how to flow data from remote sites, via a Web browser, into other programs running natively on the desktop, such as Integrated Genome Browser. Web browser development communities are constantly changing and improving their security models, essential to keeping users and their data safe in an increasingly adversarial and dangerous digital environment. Most Web pages are now loaded over encrypted channels, using HTTPS, the secure version of HTTP, and this includes BioViz *Connect*. This means that the JavaScript code responsible for interacting with IGB’s localhost endpoint is also loaded via HTTPS. However, when this code interacts with IGB via its localhost endpoint, it does so via unencrypted HTTP, because there is currently no robust way to support HTTPS for the localhost domain. The Chrome and Firefox browser allow BioViz *Connect* code to access the localhost IGB endpoint using HTTP because the communication channel is limited to the user’s own computer, presumed to be secure. The MacOS Safari Web browser does not allow it, however. This means that that BioViz *Connect’s* “View in IGB” feature fails for Safari users. We handle this by advising the user to switch to a different browser on MacOS. However, the more permissive Chrome and Firefox browsers may block this interaction in their future releases. This issue exemplifies a more general problem with connecting the desktop to the cloud. The methods used to communicate with remote computers are always changing, usually becoming more restrictive, which means that developers need to constantly test, revise, and update their software, more so perhaps than developers who create stand-alone, independent applications that rarely need to interoperate with anything other than the host computer’s operating system.

Architectures using Web-based REST APIs may help solve these problems. For example, CyVerse or BioViz *Connect* could add new endpoints that themselves support region-based retrieval of genomic data, as with the XML-based Distributed Annotation Service (Dowell et al., 2001; Jenkinson et al., 2008) and the newer JSON-based University of Santa Cruz Genome Informatics REST interface (UCSC). Rather than deliver data in new JSON or XML formats that would require modifying the client software, these new endpoints could simply stream the data in their native formats. Ideally, these new API endpoints would require minimal or no change to the client software. Another way to achieve this would be to design APIs using the facade design pattern, in which an application translates an incompatible interface to a compatible one, expanding the range of clients able to access a resource. For example, to solve the public URL problem mentioned above, developers could create a novel API that provides all the services required for accessing BAM files and their indexes, by creating and destroying secure URLs as users open and load data file resources during a session. Many variations are possible, and as cloud computing infrastructures become easier and cheaper to build upon, more bioinformatics groups will attempt even more daring and exciting innovations, amplifying their users’ ability to investigate biological systems.

Finally, we highlight aspects of the BioViz *Connect* interface and functionality that could be further developed to help users find useful tools and help developers find users for their tools. First, we note that the “View in IGB” button in the BioViz *Connect* table view occupies a column labeled “Visualization Tools,” a space where links to other visualization tools could also be added, based on the input data they accept. To make space for these other tools, we could replace the button with an IGB logo, and use tooltips to provide documentation or link to videos describing how to use the tools. Second, we could enhance BioViz *Connect* search capabilities to query MetaData tags or other file properties and attributes. Third, we could collaborate with the CyVerse team and other users to design and implement data registries, which data providers and users could use to publish, publicize, and locate data sets relevant to their work. As we hope the name suggests, BioViz *Connect* will connect researchers with data and tools, and will help tool developers connect with their intended audience, improving scientific practice for everyone.

## Conflict of Interest

*The authors declare that the research was conducted in the absence of any commercial or financial relationships that could be construed as a potential conflict of interest*.

## Author Contributions

NHF and AEL conceived of and supervised the project. KR, CK, ST, and PB planned and developed BioViz *Connect*. NHF, AEL, KR, CK, ST, PB, and CD tested and debugged BioViz *Connect*. NHF, KR, CK, and AEL wrote the draft manuscript. All authors read and approved the final manuscript.

## Funding

Research reported in this publication was supported by the National Institute of General Medical Sciences of the National Institutes of Health under award number 5R01GM121927. Funding was used to plan, design, and develop the software reported in the article.

## Abbreviations

API: Application Programming Interface;
CSS: Cascading Style Sheets;
HTML: HyperText Markup Language;
IGB: Integrated Genome Browser;
REST: REpresentational State Transfer

## Acknowledgements

We thank Paul Sarando, Sarah Roberts, Sriram Srinivasan, Ian McEwen, Ramona Walls, and Reetu Tuteja for their assistance with the Terrain API and publishing CyVerse apps, which was made possible through CyVerse’s External Collaborative Partnership program.

## Project links

https://bitbucket.org/lorainelab/bioviz-connect, https://bitbucket.org/lorainelab/bioviz-connect-playbooks

## Operating system(s)

Linux-based systems for the BioViz *Connect* server. BioViz Connect web app tested on Ubuntu, Windows 10, MacOS Mojave operating systems and Chrome, Firefox, and Edge web browsers.

## Programming language

HTML, JavaScript, CSS, and python3

## Other requirements

A free CyVerse account is required to login: https://user.cyverse.org/register. Integrated Genome Browser 9.1.4 or greater required for visualizing data using BioViz *Connect*.

## License

Common Public License Version 1.0 Any restrictions to use by non-academics: No

## Notes

https://bioviz.org

https://bitbucket.org/lorainelab/bioviz-connect-playbooks

https://bitbucket.org/lorainelab/bioviz-connect

### Competing Interest Statement

The authors have declared no competing interest.

### Summary of Updates

This new version contains a Discussion section describing the potential impact and significance of the work.

